# HepI and OpsX are functionally coupled but evolutionarily asymmetric heptosyltransferase variants: ecological transitions and operon modularization drive divergent constraints and flexibility

**DOI:** 10.64898/2026.06.09.731171

**Authors:** Aayatti Mallick Gupta, Philip Arevalo, Erika Anne Taylor

**Affiliations:** Wesleyan University, 52 Lawn Ave., Middletown, Connecticut, 06459, U.S.A

**Keywords:** Lipopolysaccharide biosynthesis, Heptosyltransferase I (HepI/rfaC/waaC), OpsX, Gram-negative bacteria, Molecular evolution, Horizontal gene transfer, Purifying selection, Operon architecture

## Abstract

Lipopolysaccharide (LPS) inner-core biosynthesis is classically initiated by a heptosyltransferase enzyme most commonly Heptosyltranferase I (HepI), a conserved WaaC-like enzyme. An alternative heptosyltranferase variant, OpsX, occurs alone in a subset of Gram-negative bacteria and co-exist with WaaC-like enzyme within the same genome of other organisms, raising questions about the origin of these two vairants and their functional partitioning. Here, we present a comparative evolutionary analysis of HepI (K02841) and OpsX (K12982) across Gram-negative bacteria to resolve their functional coupling and divergence. Selection analyses reveal a consistent evolutionary asymmetry, with OpsX exhibiting elevated ω values relative to HepI across global datasets and within genomes encoding both systems. Residue-level analyses indicate conserved catalytic cores in both enzymes, but a broader distribution of relaxed constraints in OpsX, suggesting differential partitioning of functional pressure. HepI has undergone intensified purifying selection in host-associated lineages, whereas OpsX shows no corresponding shift, indicating distinct responses to ecological context. Gene–species tree reconciliation further reveals contrasting horizontal gene transfer (HGT) architectures: HepI displays an ecologically structured network enriched in pathogen- and opportunist-associated lineages, with recurrent hub-mediated exchanges and deeper lineage-integrated events, whereas OpsX exhibits a diffuse transfer regime dominated by non-pathogenic taxa and primarily recent terminal acquisitions. These differences persist in genomes co-encoding both systems, where HepI transfer signal remains strongly associated with lifestyle, while OpsX is largely uncoupled from ecological structure. Analysis of operon architecture reveals pathway partitioning between the two genes: HepI is embedded in a conserved downstream operon linked to glycosyltransferase-mediated core assembly, whereas OpsX occurs in a more variable context enriched for upstream ADP-heptose precursor biosynthesis genes. In dual-system genomes, HepI is reduced to a minimal downstream module while OpsX retains upstream functions, indicating coordinated operon modularization. Together, HepI and OpsX form a functionally coupled but evolutionarily asymmetric system shaped by ecological transitions and genomic reorganization.

**Highlights:**

- HepI and OpsX represent functionally coupled but evolutionarily asymmetric LPS inner-core biosynthesis systems across Gram-negative bacteria.
- OpsX shows relaxed selective constraint and a diffuse horizontal gene transfer pattern, whereas HepI is under stronger purifying selection and ecologically structured transfer.
- Operon organization reveals pathway modularization, with HepI embedded in conserved downstream assembly modules and OpsX retaining upstream precursor-associated flexibility.

## Introduction

The outer membrane of Gram-negative bacteria is fortified by lipopolysaccharide (LPS), a tripartite glycolipid composed of lipid A, the core oligosaccharide, and, in many taxa, the O-antigen (Raetz and Whitfield 2002; Whitfield and Trent 2014; Whitfield et al. 2020). Beyond its structural role, LPS is a major determinant of membrane permeability, environmental resilience, immune recognition, and host colonization. Modifications in the LPS core can profoundly alter susceptibility to complementation, antimicrobial peptides, bile salts, and antibiotics, while also reshaping host–pathogen interactions (Raetz and Whitfield 2002; Whitfield and Trent 2014; Needham and Trent 2013). Because the inner core forms the biochemical bridge between lipid A and the distal glycan architecture, enzymes acting at this stage often experience strong functional constraint while simultaneously serving as key substrates for adaptive diversification. Among these enzymes, heptosyltransferase I (HepI; canonical WaaC; KEGG K02841) catalyzes the first committed heptose transfer to the nascent lipid A–3-deoxy-D-*manno*-oct-2-ulosonic acid (Kdo) acceptor and represents one of the earliest conserved steps in inner-core assembly (Gronow et al. 2000). In classical enterobacterial systems such as *Escherichia coli*, *Salmonella enterica*, *Klebsiella pneumoniae,* and related taxa, WaaC transfers L-glycero-D-*manno*-heptose to a lipid A structure containing two Kdo residues, thereby establishing the conserved heptose backbone of the LPS inner core (Gronow et al. 2000; Sirisena et al. 1992; Noah et al. 2001). Loss of this reaction produces “deep-rough” phenotypes with severely truncated LPS and destabilized outer membranes, highlighting the essential structural role of HepI-mediated heptosylation (Heinrichs et al. 1998). Accordingly, WaaC homologs were historically viewed as a broadly conserved and functionally uniform enzyme family across Gram-negative bacteria (Sirisena et al. 1992; Stojiljkovic et al. 1997; Gronow et al. 2000; Raetz and Whitfield 2002).

This paradigm was challenged by studies in *Haemophilus influenzae*, where no canonical *waaC* ortholog was identified, but instead a homolog of *opsX* (Gronow et al. 2000)—originally described in *Xanthomonas campestris* as a gene implicated in LPS biosynthesis—was detected and subsequently assigned as an alternative first heptosyltransferase system (Kingsley et al. 1993; Hood et al. 1996; Cody et al. 2000). Biochemical characterization demonstrated that OpsX encodes a structurally distinct heptosyl-I like transferase that transfers the first heptose only to phosphorylated Kdo (Kdo-P), rather than to the classical di-Kdo acceptor, revealing a mechanistically divergent route for inner-core initiation (Gronow et al. 2000). Independent genetic evidence from *Haemophilus parasuis*, an important porcine pathogen and the etiological agent of Glässer’s disease, further reinforced this role, as deletion of *opsX* produced severely truncated lipooligosaccharide (LOS) structures consistent with disruption of early inner-core heptosylation ((Zhou et al. 2015; Xu et al. 2016). Together, these findings established OpsX as a novel class of heptosyltransferase I enzyme and suggested that apparently conserved LPS assembly pathways can evolve alternative catalytic solutions through substrate rewiring rather than wholesale pathway replacement. Subsequent comparative work proposed that some lineages may harbor both WaaC-like and OpsX-like systems, but these observations remained limited to a few species (Logan et al. 2006) and lacked a broader evolutionary framework. A second major insight emerged from *Pasteurella multocida*, where two simultaneously expressed heptosyl-1 transferase enzymes generate alternative inner-core glycoforms (Harper et al. 2007). In this species, HptA (OpsX-like) specifically recognizes lipid A–Kdo₁-P, whereas HptB (WaaC-like) acts on lipid A–Kdo₁–Kdo₂. Together with the Kdo kinase KdkA and bifunctional Kdo transferase KdtA, these enzymes enable parallel biosynthesis of distinct LPS inner-core architectures within the same cell population. Related observations in *Haemophilus parainfluenzae* further reinforce this view (Young and Hood 2013). Unlike the highly phase-variable LPS repertoire of *Haemophilus influenzae, H. parainfluenzae* exhibits a more restricted set of outer-core genes and reduced phase variation, yet retains both *opsX* and *waaC* homologs. This combination has been interpreted as evidence for latent inner-core flexibility coupled with constrained outer-core diversification (Humphries and High 2002). Collectively, these lineage-specific studies imply that HepI system composition may influence the balance between structural robustness and adaptive plasticity, but they do not reveal how widespread such strategies are across Gram-negative bacteria or whether they are associated with particular ecological transitions.

Despite decades of biochemical and structural work on LPS biosynthesis, the macroevolutionary history of alternative heptosyltransferase I systems remains unresolved. It is unknown how often OpsX arose relative to canonical WaaC, whether dual heptosyltransferase I architectures represent transient intermediates or stable genomic states, and how these patterns relate to host association, opportunism, pathogenicity, or free-living environmental lifestyles. More fundamentally, because HepI and OpsX catalyze sugar addition in analogous pathway positions but differ in substrate specificity and genomic context, they offer an opportunity to test how functional coupling can coexist with unequal evolutionary constraint. If one variant primarily preserves indispensable core structure while the other mediates pathway flexibility, then asymmetric selection, gene turnover, and operon dynamics should leave detectable comparative signatures. Our previous structural and evolutionary analysis of heptosyltransferase I variants in Proteobacteria provided evidence for strong conservation of catalytic architecture despite substantial sequence divergence, suggesting that functional constraints dominate over sequence-level variability (Gupta et al. 2025). Here, we presented the first large-scale comparative evolutionary analysis of HepI (K02841) and OpsX (K12982) across nearly two thousand Gram-negative bacterial species. By integrating phylogenetic distribution, ecological metadata, operon organization, gene neighborhood architecture, and codon-level evolutionary analyses, we moved beyond genus-specific case studies to a genome-wide view of early LPS core evolution. We specifically distinguished genomes encoding HepI alone, OpsX alone, or both, thereby resolving an underappreciated dimension of LPS biosynthetic diversity: the stable coexistence of alternative first-heptosyltransferase systems within the same genomic background. Such study provided a comprehensive evolutionary framework for two different glycoforms clarify earlier species-level observations where the predominant first heptosyltransferase corresponded to HepI rather than OpsX, and demonstrated how pathway-essential enzymes can diversify through modularization without loss of core function.

## Results

### Comparative dataset structure for HepI and OpsX across lifestyles

A comprehensive dataset of the two variants of heptosyltransferase I - HepI and OpsX was constructed by systematically querying the Annotree (version r214) (Mendler et al. 2019) database using KEGG Orthology identifiers K02841 and K12982, respectively. The initial retrieval encompassed all annotated hits across available bacterial genomes, followed by stringent curation to ensure phylogenomic reliability and biological relevance. Uncultured metagenome-assembled genomes (MAGs), incomplete fragments, and low-quality assemblies were excluded to minimize annotation uncertainty. In addition, type strains were removed to avoid phylogenetic bias, and redundant entries were collapsed by retaining a single representative per species, thereby generating a non-redundant, species-level dataset suitable for comparative evolutionary inference.

For HepI (K02841), the final curated dataset comprised 1,963 unique species, spanning 645 genera across 21 bacterial phyla, representing a broad phylogenetic distribution within Gram-negative lineages and is referred to as the global HepI dataset throughout this study. The majority of sequences were concentrated within *Pseudomonadota* (n = 1,743 species), followed by contributions from *Bacillota_C* (107), *Planctomycetota* (99), *Campylobacterota* (59), *Desulfobacterota* (52), and multiple additional low-frequency phyla including *Verrucomicrobiota*, *Chlamydiota*, *Nitrospirota*, and others, collectively capturing deep evolutionary diversity across bacterial lifestyles and habitats. In contrast, the OpsX (K12982) dataset exhibited a more restricted phylogenetic distribution, comprising 977 unique species, 170 genera, and 6 phyla forming global OpsX dataset for this study, with a strong dominance of *Pseudomonadota* (n = 977 species). Minor representation was observed in *Desulfobacterota_F*, *Desulfobacterota*, *Chrysiogenota*, *Verrucomicrobiota*, and *Gemmatimonadota*, indicating a comparatively narrower evolutionary breadth relative to HepI. At the genus level, both datasets revealed strong enrichment in ecologically and clinically relevant Gram-negative lineages. In the HepI dataset, broad genus-level diversity was observed across hundreds of taxa distributed sparsely across multiple phyla, whereas the OpsX dataset showed strong overrepresentation in marine and opportunistic bacterial genera, including *Vibrio* (134), *Halomonas* (87), *Shewanella* (79), *Marinobacter* (46), *Pseudoalteromonas* (41), *Aeromonas* (32), and *Legionella* (31), among others.

Notably, comparative analysis of the curated datasets identified 69 species-level taxa in which both KEGG orthologs were independently detected. These co-occurring taxa were distributed primarily within major Gram-negative genera, including *Actinobacillus*, *Alteromonas*, *Mannheimia*, *Pasteurella*, *Rodentibacter*, *Haemophilus*, *Pseudoalteromonas*, and *Geomonas*, among others. These shared lineages were predominantly affiliated with *Pseudomonadota*, with additional representation from ecologically and clinically relevant clades such as Pasteurellaceae and marine Gammaproteobacteria. Importantly, these taxa were not merged into a single dataset but were treated as a distinct coexistence subset, as they independently encode both HepI and OpsX homologs, thereby providing a biologically meaningful framework to investigate potential functional redundancy, lineage-specific retention, and differential evolutionary trajectories of LPS core biosynthesis pathways.

Lifestyle metadata were systematically derived from genome-associated records obtained from the BV-BRC database (Davis et al. 2020), where each species was classified into ecological categories—Pathogen, Opportunist, or Non-pathogen—based on curated and aggregated annotations of host association, isolation source, and clinical relevance. These lifestyle annotations were subsequently mapped onto the global HepI, global OpsX, and coexisting datasets to enable a unified comparative framework across evolutionary and ecological dimensions. This integration facilitated systematic assessment of the distribution of HepI and OpsX homologs across distinct ecological niches, allowing evaluation of potential associations between gene presence, lineage diversification, and pathogenic potential within Gram-negative bacterial systems. In the global HepI dataset (n = 2,192), lifestyle composition was dominated by non-pathogens (1,323; 60.36%), followed by pathogens (417; 19.02%), opportunists (118; 5.38%), and unclassified taxa (334; 15.24%). The global OpsX dataset (n = 989) showed a similar distribution with a higher proportion of non-pathogens (688; 69.57%), followed by pathogens (114; 11.53%), opportunists (56; 5.66%), and unknowns (131; 13.25%). The coexisting dataset (n = 69) likewise exhibited enrichment of non-pathogenic lineages (46; 66.67%), with pathogens (9; 13.04%), opportunists (6; 8.70%), and unclassified taxa (8; 11.59%) comprising the remainder. Collectively, this curated dataset provides a deeply sampled, non-redundant, and phylogenetically balanced representation of Gram-negative bacterial diversity for HepI and OpsX homologs. The complete list of taxa and associated metadata is provided in Supplementary Tables S1–S3 (Global HepI, Global OpsX, and coexisting datasets, respectively) as machine-readable CSV files.

### Selection analysis reveals differential constraint between HepI and OpsX

To assess baseline selective pressures acting on both genes, global dN/dS (ω) values were estimated across the full dataset. In all cases, ω remained well below 1, consistent with pervasive purifying selection. *hepI* exhibited a mean ω of 0.0905 (SD = 0.0182), while *opsX* showed a slightly higher mean ω of 0.0984 (SD = 0.0121). Although the absolute difference is modest, the shift toward higher ω in *opsX* was consistent (Table S4), indicating relatively reduced selective constraint compared to *hepI*. The narrower dispersion of ω values in *opsX*, as reflected by its lower standard deviation, suggests more uniform selective pressure across the dataset, while *hepI* shows slightly greater variability. Confidence intervals for both genes remained far below neutrality (ω = 1), providing no evidence for gene-wide positive selection. Taken together, these results support a scenario in which both genes are strongly constrained, but *opsX* evolves under comparatively relaxed purifying selection relative to *hepI*.

To resolve site-specific evolutionary patterns, residue-wise selection profiles were inferred using FUBAR and mapped onto functionally defined regions of both proteins. FUBAR estimates site-wise ω under a model of pervasive selection and is therefore particularly suited to identifying consistent signatures of purifying constraint across alignments, although it is not designed to detect episodic or lineage-specific positive selection. Accordingly, the inferred ω values are interpreted here as relative indicators of constraint across sites rather than absolute measures of selection intensity. Across both datasets, the majority of residues exhibited ω < 1, consistent with pervasive purifying selection at the site level. In HepI, strong purifying selection is evident at key functional regions, particularly the N-terminal acceptor motifs (positions 10–14, 60–64, and 120–124), previously recognized as critical for ligand interaction and pocket architecture (Grizot et al., 2006, Gupta et al., 2025), as well as at residues within the C-terminal catalytic region (Grizot et al., 2006) (positions 188, 192, and 241). Motif-wise comparison of mean ω values shows that both HepI and OpsX remain under purifying selection across all three N-terminal acceptor motifs, but with subtle differences between them (Fig. 1a). In Motif 10–14, OpsX exhibits a slightly lower ω than HepI, indicating comparable or marginally stronger constraint at this region. In contrast, Motifs 60–64 and 120–124 show higher ω values in OpsX relative to HepI, suggesting a modest reduction in constraint at these functionally important sites, although values remain well below neutrality. These differences indicate that while the overall functional integrity of the acceptor motifs is preserved in both proteins, OpsX displays a slight shift toward increased evolutionary flexibility in specific motifs. A similar pattern is observed at key C-terminal catalytic residues (Fig. 1b). At positions 188, 192, and 241, OpsX consistently shows higher ω values compared to HepI, with the difference being most pronounced at residue 192, where HepI is highly constrained while OpsX exhibits a marked increase in ω. Residue 241 also shows a noticeable elevation in OpsX, whereas residue 188 displays a moderate increase. In contrast, residue 262 shows a weaker deviation from the overall trend, with HepI exhibiting a higher ω than OpsX, suggesting relatively stronger constraint in OpsX without a sharp site-specific shift. Overall, these comparisons indicate that although both proteins maintain conserved functional regions, OpsX shows slightly elevated ω at several key motif and catalytic residues, reflecting modest differences in site-specific evolutionary constraint rather than a fundamental shift in selective regime.

**Figure 1.**
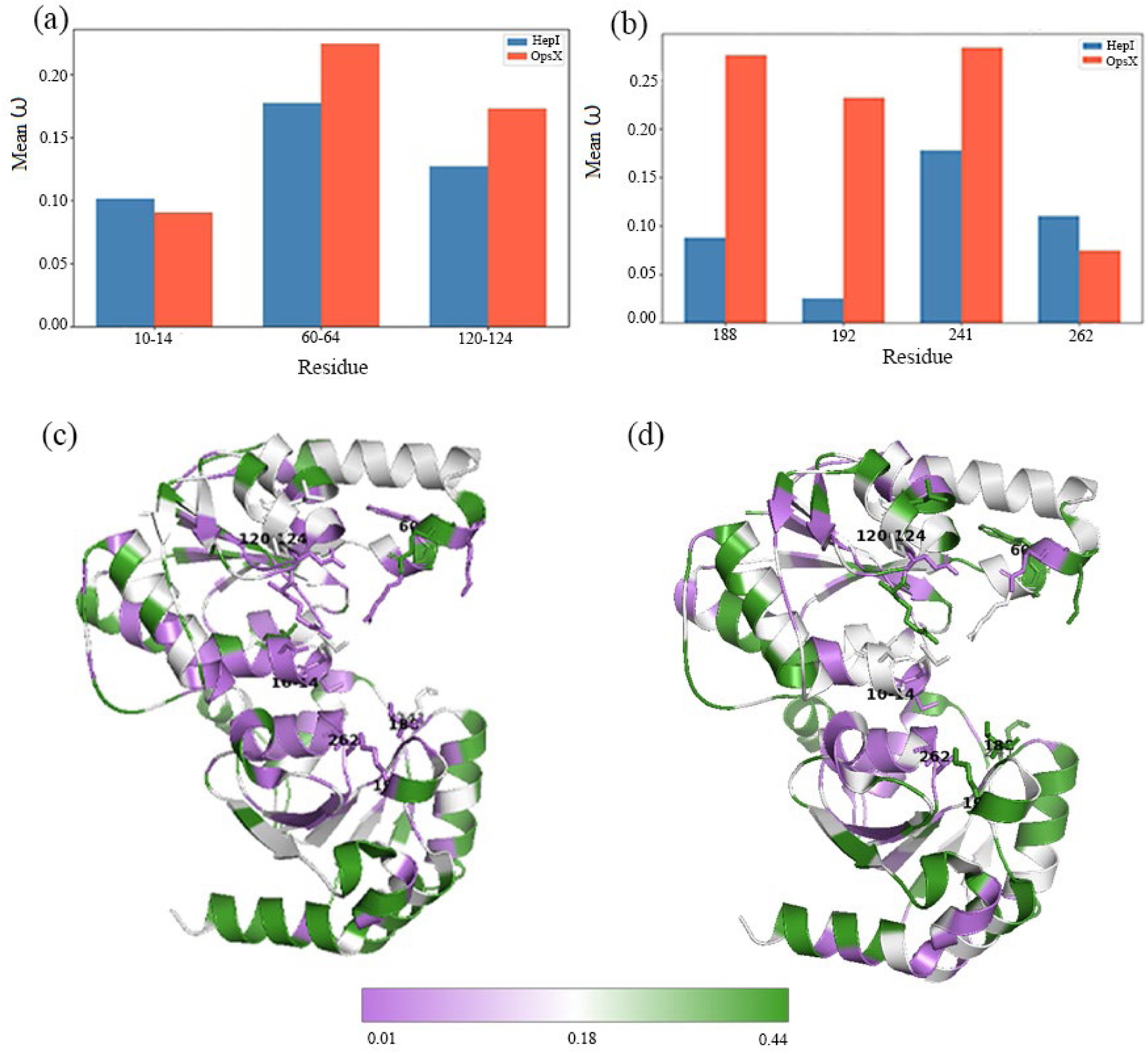
Residue-wise ω and structural evolutionary constraint in HepI and OpsX. (a) Motif-wise mean ω comparison across conserved N-terminal acceptor motifs (10–14, 60–64, 120–124) in HepI (steel blue) and OpsX (tomato red). Both proteins are under purifying selection, with OpsX showing slight relaxation in motifs 60–64 and 120–124. (b) Mean ω at key C-terminal catalytic residues (188, 192, 241, 262). OpsX shows elevated ω at most sites 188, 192 and 241, while residue 262 shows the reverse pattern; all remain under purifying selection. (c) Residue-wise ω mapped onto HepI structure (PDB: 2GT1) shows strong constraint in core and catalytic regions, with relaxed constraint at solvent-exposed sites. (d) Projection of OpsX ω onto HepI structure reveals conserved core constraint but expanded relaxed regions in surface-exposed loops, spatially segregated from the catalytic center.

Structural mapping of residue-wise mean ω values onto the HepI crystal structure (PDB ID: 2GT1) supports these trends. In HepI (Fig. 1c), residues under strong purifying selection (ω<0.3) form a tightly clustered network within the protein core and around the catalytic pocket, including the N-terminal acceptor motifs and key C-terminal catalytic residues described above. This pattern is consistent with the need to preserve both structural integrity and catalytic function. In contrast, residues with comparatively higher ω in HepI are primarily located on solvent-exposed loops and peripheral helices, indicating regions of reduced constraint that are more permissive to sequence variation. When OpsX-derived ω values are projected onto the same HepI structural framework (Fig. 1d), a) broadly similar spatial organization of constraint is observed. Core and active-site regions remain strongly conserved, reinforcing the preservation of the underlying enzymatic scaffold across paralogs. However, relative to HepI, OpsX displays a modest expansion of higher-ω residues, particularly across surface-exposed regions and flexible loop elements. These differences are consistent with the elevated ω values observed at specific motifs and catalytic positions in OpsX, although the increases remain moderate and below neutrality. Importantly, the regions exhibiting elevated ω in OpsX remain largely spatially segregated from the catalytic center defined in HepI, indicating that divergence between the paralogs is constrained to peripheral structural elements rather than the functional core. Thus, while HepI exhibits a more tightly constrained profile overall, OpsX shows a subtle redistribution of constraint, with increased evolutionary flexibility in surface-accessible regions. Collectively, these results support a model in which both proteins maintain conserved catalytic architecture, while OpsX accommodates limited divergence that may facilitate alterations in interaction interfaces or functional fine-tuning without compromising enzymatic activity.

Pairwise comparisons of ω between HepI and OpsX within species that possess both homologs further reinforce these trends. Across 69 bacterial species encoding both HepI and OpsX, OpsX exhibits consistently elevated ω relative to HepI, with 54 species (∼78%) showing higher ω in OpsX and only 15 species (∼22%) showing the opposite trend. The median difference (Δω = OpsX − HepI) of 0.1061 indicates a modest but statistically well-supported shift toward less constrained evolution in OpsX among these species (paired Wilcoxon signed-rank test: V = 1949, p = 9.405 × 10⁻⁶). This asymmetry is clearly evident across multiple visualizations. In the paired comparison plot (Fig. 2a), most species show an upward shift from HepI to OpsX, indicating consistently higher ω values for OpsX within matched genomes. The distribution of log₁₀-transformed ω values (Fig. 2b) shows a subtle rightward shift for OpsX relative to HepI, consistent with its higher median evolutionary rate. Similarly, in the scatter plot (Fig. 2c), the majority of points lie above the diagonal (y = x), confirming that OpsX ω exceeds HepI ω in most species. The distribution of Δω values (Fig. 2d) is correspondingly skewed toward positive values, with relatively few cases where HepI evolves faster, further highlighting the directional bias in evolutionary rates. Although most species exhibit small to moderate differences (Δω ≈ 0.01–0.3), a subset of 14 species shows pronounced divergence (|Δω| > 0.3). Notably, these extreme cases are predominantly driven by elevated ω in OpsX, suggesting lineage-specific acceleration of OpsX evolution in certain taxa. Consistent with this, median ω values indicate stronger overall constraint in HepI (median ω = 0.2117) compared to OpsX (median ω = 0.3259), although both remain well below neutrality, indicating pervasive purifying selection. Together, these results demonstrate that OpsX evolves consistently faster than HepI even within the same genomic contexts, ruling out taxonomic sampling as a primary driver of the observed pattern. Instead, the systematic elevation of ω in OpsX, coupled with occasional lineage-specific accelerations, supports a model of conserved functional constraint in HepI alongside modestly increased evolutionary flexibility in OpsX.

**Figure 2.**
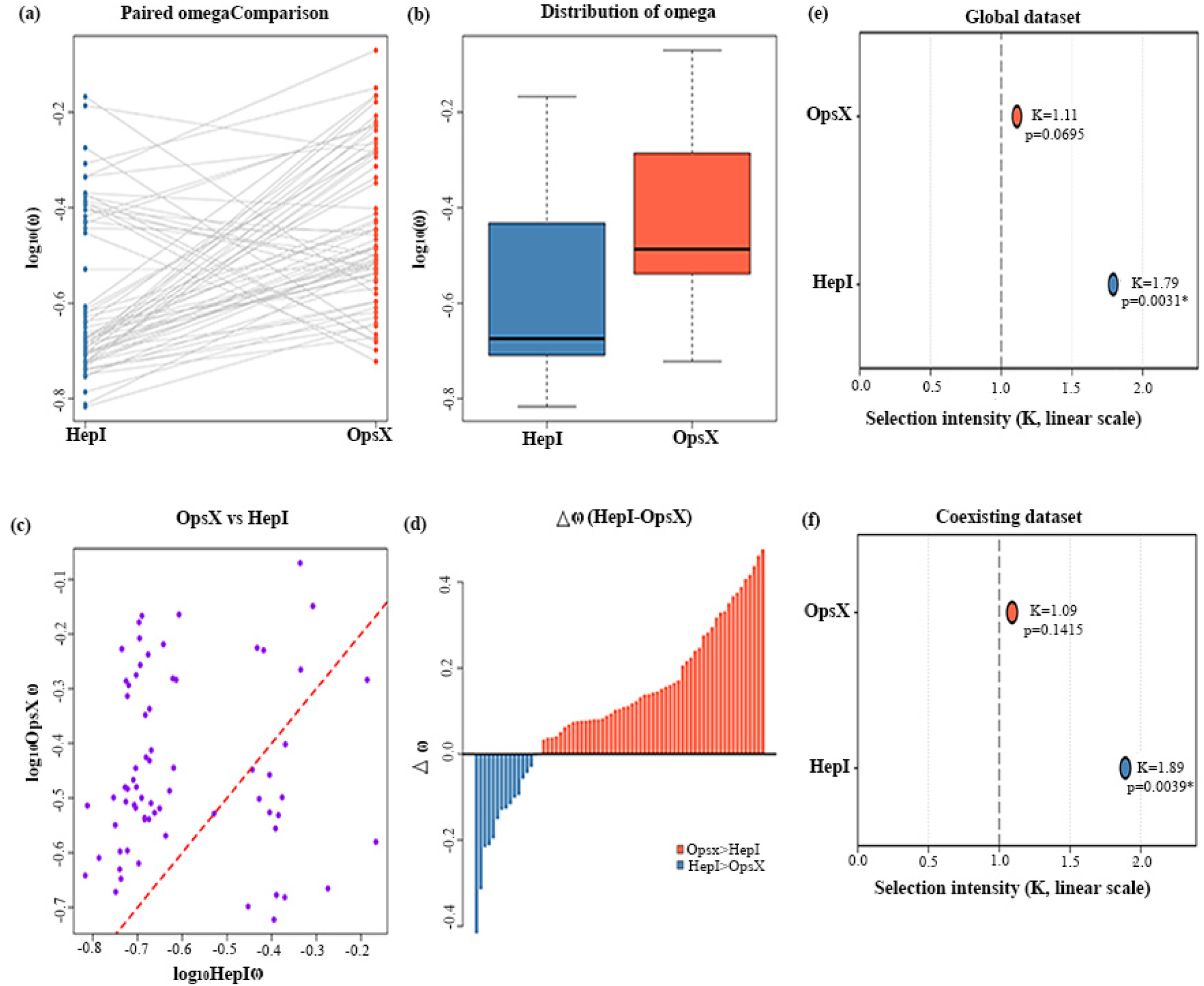
Pairwise evolutionary rate asymmetry and ecological shifts in selection between HepI and OpsX. (a) Paired ω comparisons across 69 genomes encoding both heptosyltransferase I glycoforms show a consistent upward shift from HepI to OpsX, with 54 species exhibiting higher ω in OpsX and 15 showing the reverse trend. (b) Distribution of log₁₀-transformed ω values shows a rightward shift in OpsX relative to HepI, indicating overall elevated evolutionary rates. (c) Species-wise scatter plot of ω values (OpsX vs HepI) shows most points above the y = x diagonal, confirming faster evolution of OpsX across genomes. (d) Distribution of Δω (OpsX − HepI) is skewed toward positive values (median Δω = 0.1061; Wilcoxon signed-rank test V = 1949, p = 9.405 × 10⁻⁶), with a subset of taxa showing strong divergence (|Δω| > 0.3), largely driven by OpsX acceleration. (e) Selection intensity (RELAX) in the global dataset shows intensified purifying selection in HepI (K = 1.79, p = 0.0031), with no deviation in OpsX. (f) Coexisting dataset follows the same pattern, with HepI showing significant intensification (K = 1.89, p = 0.0039) and OpsX remaining neutral (K = 1.09, p = 0.1415).

### Host-associated lineages exhibit intensified purifying selection in HepI relative to non-host-associated lineages, while OpsX shows no corresponding shift

Ecological effects on selection intensity were quantified using the RELAX framework, which models shifts in the distribution of dN/dS (ω) through a scaling parameter K applied to site-specific ω categories such that ω_i’_=ω_i_^K^. Under this formulation, K > 1 amplifies the separation among ω classes, intensifying both purifying (ω < 1) and positive (ω > 1) selection, whereas K < 1 compresses the distribution toward neutrality (ω = 1), indicating relaxation; K = 1 corresponds to no change relative to the background distribution. Thus, K reflects a global rescaling of selective pressures across sites rather than locus-specific effects. Statistical support was assessed using likelihood ratio tests against the null hypothesis (K = 1). Lifestyle annotations curated from BV-BRC were used to classify lineages into ecological categories and define contrasts between host-associated and non-host-associated taxa.

In the global dataset, a stringent ecological contrast was implemented by assigning pathogen lineages as the foreground and non-pathogens as the background. Given the larger and more diverse sampling this design maximizes ecological separation and minimizes ambiguity arising from intermediate lifestyles, thereby providing a conservative test for detecting shifts in selection intensity. Under this framework, HepI exhibited significant intensification of selection (K = 1.79, p = 0.0031; Fig. 2e), indicating stronger purifying constraint in host-associated lineages, whereas OpsX showed no detectable deviation from the null expectation (K ≈ 1; Fig. 2e). In the coexisting dataset, which is restricted to genomes encoding both genes and is therefore inherently reduced in size and taxonomic breadth, a broader definition of host association was adopted by combining pathogen and opportunist lineages into the foreground, with non-pathogens retained as the background. This adjustment is methodologically motivated to preserve statistical power and maintain sufficient representation of the foreground class, as enforcing a strict pathogen-only definition for foreground branches under reduced sampling would lead to imbalanced categories and decreased sensitivity of the likelihood-based test. Importantly, opportunist taxa represent intermediate but biologically relevant host-associated lifestyles, and their inclusion maintains the directionality of the ecological contrast while avoiding over-fragmentation of categories. Under this framework, the same pattern was observed (Fig. 2f): HepI again showed significant intensification of purifying selection in host-associated taxa (K = 1.89, p = 0.0039), whereas selective pressures on OpsX remained indistinguishable between host-associated and non-host-associated taxa. (K = 1.09, p = 0.1415).

The consistency of results across these two complementary contrast definitions—one stringent and one inclusive—indicates that the observed intensification for HepI and the stability for OpsX are not artifacts of foreground assignment but are robust to differences in ecological categorization and dataset composition. Together, these analyses show that selection intensity increases in host-associated lineages for HepI but remains unchanged for OpsX across both ecological frameworks.

### Phylogenetic structure across shared genomes encoding both HepI and OpsX reveals stronger vertical signal and lineage-specific divergence in HepI, with increased lineage mixing and reduced resolution in OpsX

Phylogenetic analysis of genomes encoding both HepI and OpsX revealed clear differences in tree topology and evolutionary signal between the two genes, despite being examined across an identical taxonomic background and rooted using HepIII as a common outgroup (Fig. 3a-b). The HepI phylogeny exhibited a well-resolved and structured topology, with distinct clustering of lineages and clear separation of major clades, indicative of strong vertical inheritance and lineage-specific divergence. In contrast, the OpsX tree showed a comparatively diffuse topology, characterized by increased intermixing of taxa and reduced clade resolution, suggesting weaker phylogenetic signal. These topological differences were further supported by quantitative branch length comparisons. Across both branch types, HepI displayed significantly longer branch lengths than OpsX. Internal branches were longer in HepI (mean = 0.18) compared to OpsX (mean = 0.11; Mann–Whitney U test, p = 0.02002), and a similar trend was observed for tip branches (HepI mean = 0.23; OpsX mean = 0.14; p = 0.01957). Within-gene comparisons revealed that HepI exhibited significantly longer tip branches relative to internal branches (p = 0.03851), reflecting enhanced terminal divergence, whereas OpsX showed no significant difference between branch types (p = 0.09549), consistent with a more homogenized evolutionary pattern. The heatmap summarizing mean branch lengths further illustrates these differences, with HepI consistently exhibiting higher values across both internal and tip branches compared to OpsX (SI Fig. S1a). Notably, the greater separation between internal and tip branch lengths in HepI contrasts with the relatively compressed distribution observed in OpsX. Together, these results demonstrate that HepI retains a stronger vertical evolutionary signal with pronounced lineage-specific divergence, whereas OpsX exhibits increased lineage mixing, reduced phylogenetic resolution, and overall lower divergence within the same set of coexisting taxa.

**Figure 3.**
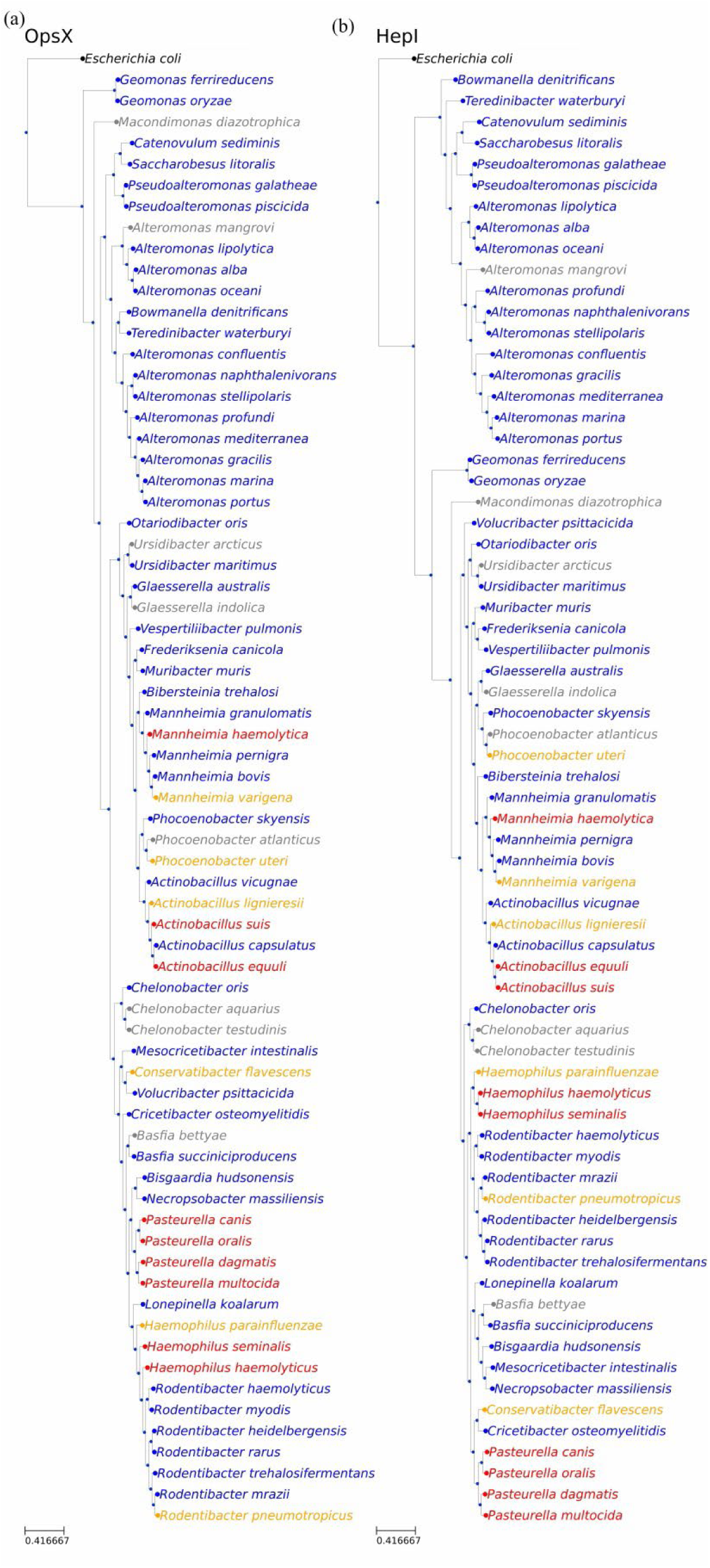
Contrasting phylogenetic structure of HepI and OpsX in coexisting taxa. Maximum-likelihood phylogenies of HepI (a) and OpsX (b) from genomes co-encoding both genes, rooted using HepIII. The HepI tree exhibits strong topological structure with well-resolved, lineage-specific clades consistent with vertical inheritance. In contrast, the OpsX phylogeny shows reduced resolution and increased lineage intermixing, indicating a weaker phylogenetic signal and greater disruption of vertical structure across the same taxonomic background.

### Reconciliation analysis reveals host-associated structuring of horizontal transfer in HepI, with diffuse transfer dynamics in OpsX

Gene–species tree reconciliation under a duplication–transfer–loss (DTL) framework (RANGER-DTL; 50 iterations with >60% event support) revealed fundamentally distinct horizontal gene transfer (HGT) architectures for *hepI* and *opsX* across both global and coexisting datasets. In the global dataset, *hepI* exhibited a highly connected and non-random transfer network, with pronounced clustering of events and the emergence of high-degree “super-spreader” hubs (Fig. 4a-c. These hubs were disproportionately composed of pathogenic and opportunistic taxa, which functioned both as donors and recipients, indicating bidirectional and recurrent transfer within ecologically interactive lineages. Consistently, ecological transfer matrices showed strong enrichment of pathogen- and opportunist-associated exchanges (Fig. 4d), supporting a model of ecologically structured HGT. In contrast, *opsX* displayed a diffuse and weakly organized network (Fig. 4e-g), with sparse backbone transfers, reduced hub centrality, and a predominance of non-pathogen lineages. Ecological bias was minimal, with transfers largely occurring among non-pathogens and lacking clear directionality (Fig. 4h), indicating a more background-driven or stochastic transfer regime. Network based analysis of HGT further supported ecological structuring in *hepI* where pathogen and opportunist taxa were prominently represented among top transfer events and donar hubs. In contrast*, opsX* exhibits a diffuse network dominated by non-pathogen lineages, with limited involvement of pathogenic taxa. Ecological flow matrices confirm that while transfer among non-pathogens occur broadly, high-weight and hub-mediated transfers in *hepI* are enriched in ecologically relevant lineages. These results indicate that HGT in hepI is driven by ecological selection, whereas *opsX* evolves under a more neutral or constrained regime.

**Figure 4.**
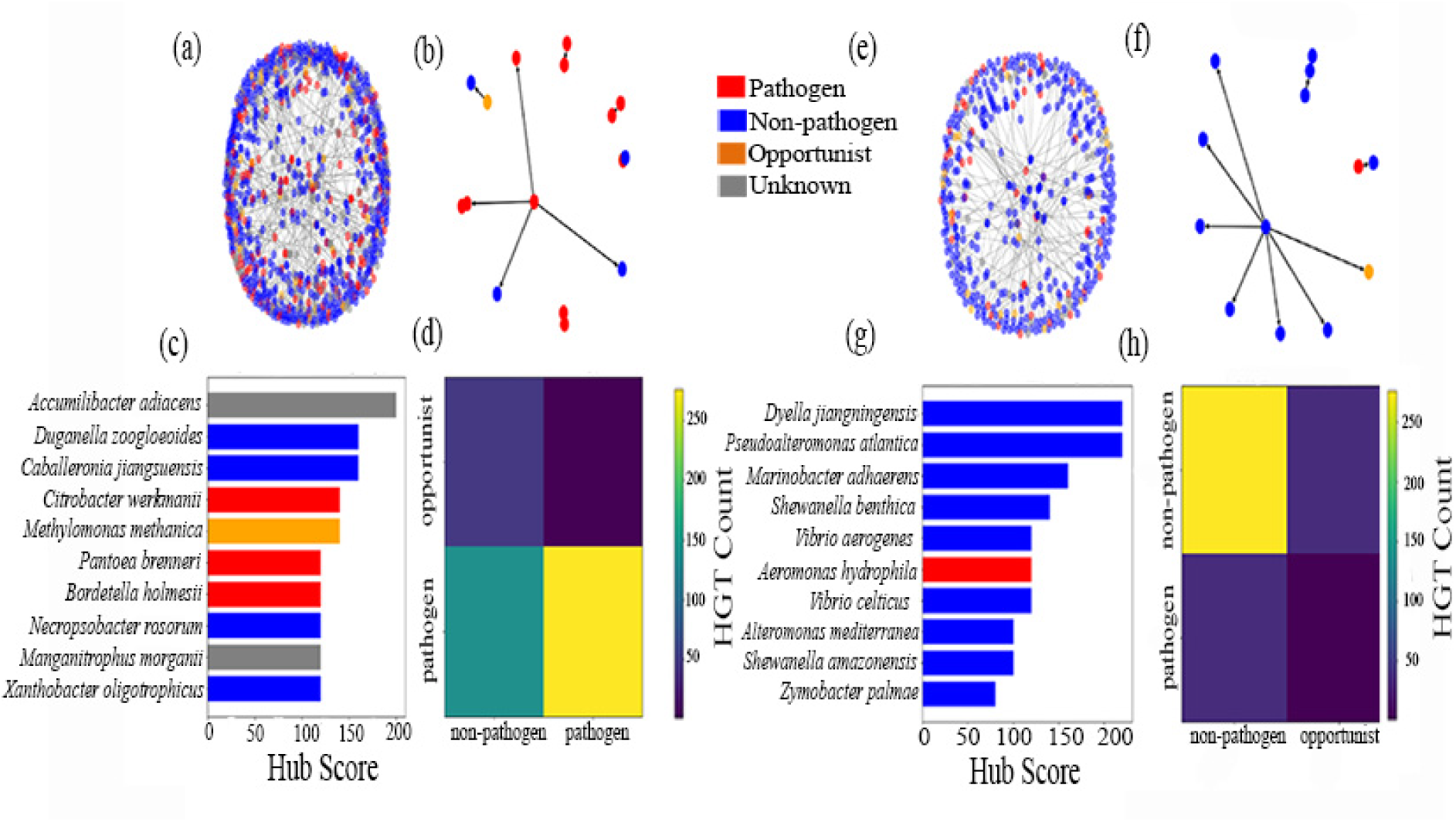
Contrasting horizontal gene transfer architectures of HepI and OpsX inferred from gene–species tree reconciliation. (a–c) RANGER-DTL reconciliation of HepI (global dataset; ≥60% event support across 50 iterations) reveals a highly structured HGT network with dense connectivity and prominent high-degree “super-spreader” hubs. These hubs are enriched in pathogen- and opportunist-associated lineages acting as both donors and recipients, indicating recurrent, bidirectional, and ecologically structured transfer. (d) Ecological transfer matrix for HepI shows strong enrichment of exchanges among pathogen and opportunist lineages, consistent with non-random, ecology-driven HGT. (e–g) OpsX reconciliation network shows a sparse, weakly connected topology with reduced hub centrality and limited clustering of transfer events. Transfers are primarily distributed among non-pathogenic taxa, indicating a diffuse and less structured HGT regime. (h) Ecological transfer matrix for OpsX shows minimal ecological bias, with broadly distributed transfers and weak signal of lineage-specific enrichment, consistent with a more stochastic or background-driven transfer process.

These patterns were also observed in the coexisting dataset, where similar counts of inferred transfer events concealed pronounced differences in their evolutionary organization. Phylogenetically controlled regression (PGLS) revealed a strong and significant association between lifestyle and HGT signal in hepI (β = 3.88, p < 10−10), which was further supported by phylogenetic permutation (p=0.014), indicating a non-random, ecology driven distribution of HGT events. In contrast, *opsX* showed no significant association (β = 0.57, p = 0.51), suggesting that HGT is not structured by ecological lifestyle (SI Fig. S2_a-b). Despite co-occurance of both genes within the same taxa, these results highlight a striking asymmetry in evolutionary dynamics, with *hepI* ecology linked transfer patterns, whereas *opsX* remains unbiased*. hepI* transfers were enriched along internal branches (∼55%; SI Fig. S2_c), consistent with deeper, lineage-integrated events that are retained and propagated within clades, whereas *opsX* transfers were predominantly terminal (∼52%), reflecting recent, lineage-specific and likely transient acquisitions. Phylogenetic reconciliation of coexisting taxa (SI Fig. S3) further supported this distinction: *hepI* showed clustered, recurrent transfer events—particularly among high-confidence transfers highlighted by branch thickness—within ecologically coherent clades, while *opsX* transfers were scattered, non-recurrent, and phylogenetically diffuse. Notably, not all inferred transfer events are explicitly visualized in the phylogram, as those mapped to deeper ancestral nodes are hierarchically embedded within internal clades and cannot always be resolved at terminal levels. Together, these results demonstrate that *hepI* is embedded within an ecologically structured HGT network dominated by pathogen and opportunist lineages, suggesting transfer driven by ecological interaction pressures such as host association, competition, and stress adaptation, whereas *opsX* participates in a more diffuse and weakly constrained transfer landscape characteristic of background genomic flux.

### Operon architecture reveals pathway partitioning and functional decoupling

To investigate the genomic context of *hepI* and *opsX*, comparative genomics–based synteny analysis was performed to resolve gene neighborhood organization and infer operon architecture. Analysis of the global dataset revealed that *hepI* is embedded within a highly conserved operon context consistent with a functionally integrated LPS inner-core biosynthetic module, comprising genes involved in precursor processing, glycosyltransferase activity, and heptose incorporation (Fig. 5a). This arrangement encodes a functionally coherent module associated with lipopolysaccharide (LPS) inner-core biosynthesis, integrating genes involved in precursor processing (*coaD*), glycosyltransfer activity (GT2, *hepI*, *hepII*), and heptose incorporation and modification (*waaA*, *rfaD*). The strong conservation of gene order suggests tight genomic coupling of downstream LPS assembly functions. The strong conservation of gene order suggests tight coupling of downstream assembly steps and sustained selective constraint on maintaining operon integrity. In contrast, global analysis of *opsX* revealed its consistent association with a distinct operon encoding enzymes required for ADP-heptose biosynthesis, representing upstream precursor-generating functions (Fig. 5b). Together, these observations indicate a functional partitioning of the LPS biosynthetic pathway, wherein *opsX* is associated with upstream precursor biosynthesis, while *hepI* retains downstream glycosyltransferase-mediated assembly functions. This separation is most evident in genomes where both genes coexist, suggesting coordinated but spatially segregated pathway organization.

**Figure 5.**
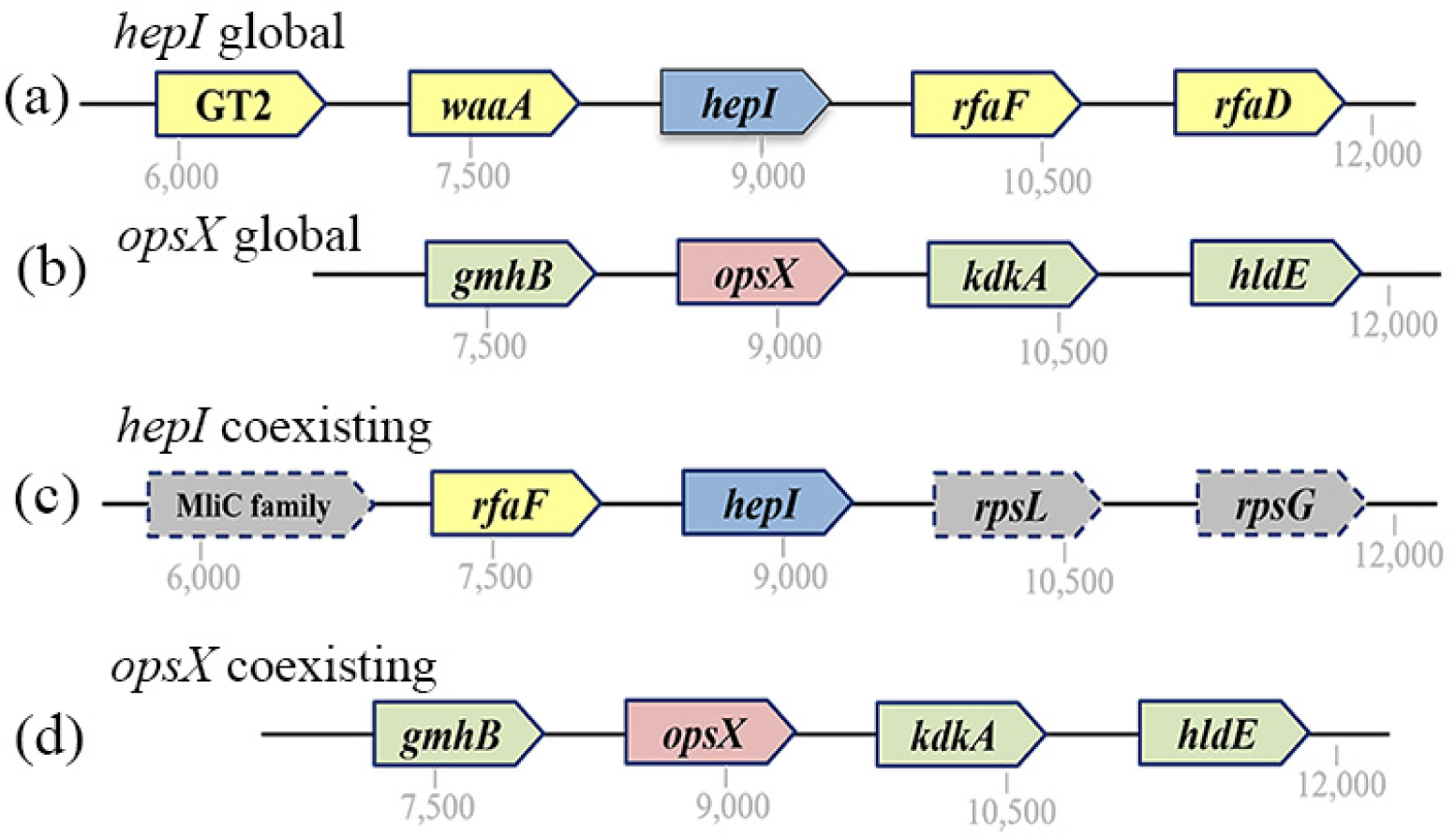
Operon architecture and genomic context of HepI and OpsX across global and coexisting datasets. (a–b) In global genomes, *hepI* (red) occurs within a conserved operon enriched in LPS inner-core biosynthesis genes (blue), whereas *opsX* (red) is embedded in a distinct upstream biosynthetic operon (olive green), indicating functional partitioning of pathway modules. (c–d) In coexisting genomes, the *hepI* operon is truncated, retaining only core downstream components (blue) with loss of upstream genes (grey, dashed), while *opsX* remains in a conserved upstream operon (olive green), maintaining precursor biosynthesis functions.

Focusing on genomes co-encoding *hepI* and *opsX* revealed a marked reorganization of these otherwise distinct modules. In such genomes, the *hepI* locus is reduced to a minimal configuration retaining core glycosyltransferase genes (*rfaC*, *waaF*) while consistently lacking upstream components such as *waaA* and *rfaD* (Fig. 5c). Concurrently, the *opsX*-associated operon remains conserved and continues to encode upstream heptose biosynthesis functions (Fig. 5d). This contrast highlights a systematic architectural shift under shared genomic contexts, wherein the canonical *hepI* operon is truncated while the *opsX* operon is retained intact. This complementary organization supports a model of functional partitioning, in which *opsX* is linked to precursor biosynthesis and *hepI* retains downstream LPS assembly functions. Within this framework, once *opsX*-associated modules ensure a stable precursor supply, upstream components of the canonical *hepI* operon become functionally dispensable. Consistent with this, genes such as *rfaD* and other upstream metabolic elements are selectively absent from the reduced *hepI* locus. Notably, these genes are not uniformly lost but instead show differential genomic repositioning: *waaA* is frequently located in proximity to the *opsX* cluster, suggesting functional reassociation, whereas *rfaD* exhibits variable genomic distribution, consistent with relaxed positional constraint.

## Discussion

The coexistence of two distinct heptosyltransferase I variants within the same Gram-negative taxa has been reported only in a limited number of cases and largely within narrow phylogenetic contexts. Early functional and biochemical studies in *Haemophilus influenzae* identified OpsX (Gronow et al. 2000) as a structurally and functionally distinct class of HepI capable of transferring heptose exclusively to phosphorylated Kdo, thereby expanding the known mechanistic diversity of LPS inner core assembly. Similarly, in *Pasteurella multocida*, the simultaneous presence of two acceptor-specific heptosyltransferases (HptA/OpsX-like and HptB/WaaC-like) enables parallel synthesis of alternative inner core glycoforms (Harper et al. 2007), suggesting that duplication and subsequent functional specialization of HepI enzymes can directly contribute to LPS structural heterogeneity. Additional observations in *Haemophilus parainfluenzae*, which exhibits a constrained LPS core repertoire and limited phase variation, further support the notion that variation in heptosyltransferase composition is closely linked to structural plasticity and host interaction potential (Young and Hood 2013). However, despite these isolated insights, the broader evolutionary distribution of alternative HepI systems across bacterial lineages—particularly in relation to pathogen, opportunist, and non-pathogen lifestyles, as well as host-associated versus environmentally unassociated taxa—remains largely unexplored. In this context, our current study provides the first large-scale comparative evolutionary framework of HepI (K02841) and OpsX (K12982) across Gram-negative bacteria, moving beyond genus-specific observations to a genome-wide perspective spanning nearly two thousand species. Importantly, by systematically distinguishing taxa encoding either enzyme or both variants simultaneously, we resolve an underappreciated dimension of LPS biosynthetic diversity: the stable coexistence of alternative heptosyltransferase I systems within the same genomic background. This finding extends earlier isolated reports and demonstrates that dual HepI/OpsX architectures are not rare anomalies but represent a phylogenetically structured feature enriched in specific Gram-negative lineages, particularly within Proteobacteria-associated clades.

From an ecological perspective, integrating lifestyle metadata further clarifies these patterns. Both HepI and OpsX are broadly enriched in non-pathogenic lineages, with comparatively lower representation in obligate pathogens and opportunistic taxa. This distribution challenges a strictly pathogen-centric interpretation of LPS diversification and instead suggests that the evolutionary maintenance of two heptosyltransferase I forms is more strongly associated with ecological versatility than with virulence alone. Notably, taxa encoding both glycoforms are predominantly affiliated with ecologically flexible Gram-negative genera spanning marine, host-associated, and environmentally occuring niches, implying that dual LPS biosynthetic architectures may confer selective advantages in fluctuating or multi-niche environments. Our previous structural and evolutionary analysis of HepI in Proteobacteria provided evidence for strong conservation of catalytic architecture despite substantial sequence divergence (Gupta et al. 2025), suggesting that functional constraints dominate over sequence-level variability. This study extends that framework by integrating OpsX into a unified comparative context, enabling direct evaluation of divergence, retention, and potential functional partitioning relative to HepI. Collectively, these observations suggest that HepI and OpsX are not merely redundant enzymes but may represent evolutionarily tuned solutions to distinct biochemical and ecological pressures acting on LPS core biosynthesis.

At the molecular level, selection analyses reveal consistent purifying selection acting on both enzymes, reflecting their essential roles in LPS core biosynthesis. However, HepI is under significantly stronger constraint, whereas OpsX exhibits a modest and systematic relaxation. Importantly, this difference is not driven by lineage composition but persists within shared genomic backgrounds, indicating intrinsic divergence in evolutionary constraint. Site- and structure-level analyses further show that while catalytic cores remain conserved in both proteins, OpsX tolerates localized increases in variability, particularly in peripheral and substrate-interacting regions, consistent with limited functional tuning rather than functional replacement. These patterns support a model of asymmetric evolutionary constraint following functional diversification, where HepI remains a highly constrained core biosynthetic enzyme, while OpsX retains biochemical conservation but with increased permissiveness to variation. The absence of positive selection across both systems suggests that divergence is not driven by recurrent adaptive sweeps, but rather by differential constraint and functional flexibility within a conserved catalytic framework.

The contrasting phylogenetic architectures of HepI and OpsX suggest fundamentally different modes of evolutionary history operating within the same genomic background. We hypothesize that HepI is predominantly governed by strong vertical inheritance, with lineage-specific divergence reflecting long-term co-evolution with host-associated ecological pressures and constrained functional optimization. The higher branch length separation and well-resolved topology are consistent with sustained accumulation of lineage-specific substitutions under stable functional constraint. In contrast, the diffuse topology and reduced branch structure in OpsX suggest a more dynamic evolutionary regime, potentially shaped by higher rates of lineage mixing, episodic gene turnover, or relaxed constraints on deep phylogenetic divergence. We hypothesize that OpsX is more permissive to evolutionary interchange across lineages, leading to reduced phylogenetic resolution and compressed divergence patterns. Collectively, these patterns support a model in which HepI behaves as a vertically conserved core enzyme with strong lineage fidelity, whereas OpsX reflects a more flexible and evolutionarily labile component within the shared LPS biosynthetic landscape. The contrasting reconciliation patterns indicate that horizontal gene transfer is not uniformly distributed across the two systems but is instead differentially shaped by ecological context and functional integration. We hypothesize that HepI is embedded within an ecologically “wired” transfer network, where host-associated lineages act as persistent conduits for gene exchange, enabling recurrent and non-random transfer events that are subsequently stabilized within clades. This suggests that HepI transfer is not merely stochastic acquisition but is filtered by ecological compatibility and maintained through selective retention in host-relevant backgrounds, reinforcing its role in adaptive LPS core remodeling under host-driven pressures. In contrast, the diffuse and weakly structured transfer landscape observed for OpsX suggests a fundamentally different evolutionary regime, where horizontal acquisition occurs more opportunistically and lacks strong ecological or phylogenetic reinforcement. We hypothesize that OpsX is subject to transient gene flux with limited long-term integration, reflecting either relaxed functional constraints or reduced coupling to host-associated selective environments. Collectively, these findings support a model in which HepI is actively embedded in ecologically structured horizontal exchange networks, whereas OpsX evolves within a more background-like transfer regime characterized by low specificity and weak ecological imprint.

To examine whether translational selection underlies the contrasting evolutionary patterns observed between *hepI* and *opsX*, we performed a comprehensive, multi-metric assessment of codon usage bias within genomes co-encoding both heptosyltransferase-I. By restricting the analysis to shared genomic backgrounds, we minimized confounding effects arising from lineage-specific nucleotide composition and ensured that comparisons reflect gene-specific rather than genome-wide biases. Across all evaluated metrics—GC content at synonymous third codon positions (GC3), effective number of codons (ENC), relative synonymous codon usage (RSCU), and codon adaptation index (CAI)—no statistically significant differences were detected between *hepI* and *opsX* (SI FigS4_a-d). The similarity in GC3 distributions indicates that both genes are subject to comparable underlying mutational biases. Likewise, overlapping ENC values suggest a similar degree of codon usage uniformity, arguing against differential constraints on synonymous codon choice. At the level of individual codons, RSCU-based comparisons failed to reveal any consistent or systematic shifts in codon preference between the two glycoforms. Furthermore, CAI values—used here as a proxy for translational optimization relative to highly expressed housekeeping genes—were statistically indistinguishable, indicating that both *hepI* and *opsX* are similarly adapted to the host translational machinery. Taken together, these results strongly suggest that codon usage bias and translational selection do not play a measurable role in shaping the distinct evolutionary dynamics of *hepI* and *opsX*.

The observed operon architectures indicate a clear functional compartmentalization within the LPS biosynthetic pathway, where gene order reflects division of labor rather than simple co-location of pathway components. We hypothesize that *hepI* represents a structurally constrained downstream assembly module whose conservation is maintained by the requirement for coordinated glycosyltransferase activity, whereas *opsX* is embedded in a more flexible upstream precursor-supply module that is evolutionarily and genomically more mobile. This separation suggests that pathway integrity is not maintained as a single operonic unit but instead distributed across modular genetic contexts that can be independently reorganized. In genomes where both variants coexist, the systematic reduction of the *hepI*-associated locus alongside retention of the *opsX* operon suggests a functional rerouting of upstream metabolic dependency. We hypothesize that *opsX* effectively assumes control of precursor generation, thereby decoupling *hepI* from its ancestral upstream biosynthetic inputs and allowing selective loss or relocation of previously essential genes. The differential repositioning of components such as *waaA* and *rfaD* further supports a model in which upstream biosynthetic functions are selectively reassigned or relaxed, while downstream assembly functions remain structurally constrained. These observations argue against simple redundancy, paralog replacement, or degenerative loss, and instead support adaptive operon remodeling through division-of-labor, whereby a previously contiguous pathway is reorganized into two specialized modules. Such restructuring likely confers regulatory advantages: precursor biosynthesis encoded by the *opsX* operon is metabolically costly and may be subject to environment-responsive regulation, whereas downstream LPS assembly mediated by *hepI* is structurally essential and more closely linked to basal cellular processes. The separation of these modules therefore enables independent regulation of metabolic input and structural output. Collectively, these findings indicate that the truncated *hepI* architecture observed in coexisting genomes reflects adaptive pathway decoupling and regulatory optimization, highlighting an evolutionary strategy that preserves functional output while increasing modular flexibility.

Genomic context analysis across both global and coexisting datasets reveals no consistent enrichment of mobile genetic element (MGE) signatures in the vicinity of either *hepI*- or *opsX*-associated loci. Canonical mobility-associated features—including insertion sequences, integrases/recombinases, prophage-related genes, conjugation machinery (including type IV secretion system components and VirB homologs), relaxases, and plasmid-associated replication or maintenance genes—are largely absent from the genomic neighborhoods of both systems (Frost et al. 2005; Juhas et al. 2009; Smillie et al. 2010). A small number of isolated cases with nearby MGE-associated annotations were detected in the global and coexisting contexts (SI FigS5_a-d); however, these occurrences are sporadic, non-recurrent across lineages, and represent a very minor fraction relative to the total analysis. Importantly, they do not show any consistent association with either *hepI* or *opsX* loci and do not form any discernible lineage-specific or locus-specific pattern. The overall pattern therefore does not support recent mobile element–driven acquisition. Instead, it is more consistent with long-term evolutionary remodeling of genomic neighborhoods, reflecting gradual lineage-specific reorganization rather than recent horizontal transfer events.

In the broader context of LPS evolution, these findings align with the notion that core biosynthetic enzymes can exhibit graded levels of constraint depending on their functional centrality and interaction complexity (Opiyo et al. 2010). The differential evolutionary dynamics of HepI and OpsX thus provide insight into how the two heptosyl-1 transferase enzyme variants within the same pathway can balance conservation and flexibility, enabling robustness of essential cellular processes while allowing incremental diversification. We further explored whether the coexistence of HepI and OpsX is associated with variation in downstream lipid A architecture. To address this, we implemented a gene-content–based pipeline to infer lipid A chemotypes directly from genome sequences of co-encoding taxa. This approach leverages the presence or absence of key biosynthetic and modification genes (e.g., *lpxL*, *lpxM*, *arnT*, *eptA*, *waaA*) to reconstruct probable acylation states and core modifications (Hummels 2025; Needham and Trent 2013; Simpson and Trent 2019). The resulting distribution reveals a striking and somewhat counterintuitive trend: lipid A is predominantly tetra-acylated across taxa, with the *tetra-acyl | Kdo₂ | PEtN* chemotype emerging as the most prevalent configuration (SI Fig S1b) in contrast to more complex hexa-acylated forms—traditionally associated with robust membrane stability and strong immunostimulatory potential (Raetz and Whitfield 2002; Needham and Trent 2013; Park and Lee 2013). This pattern suggests that, within the subset of genomes encoding both heptosyltransferase I variants, selective pressures may favor structurally simplified lipid A architectures, potentially as a strategy to modulate host immune recognition or adapt to specific ecological niches. The widespread presence of phosphoethanolamine (PEtN) modification further supports this interpretation, as such modifications are known to influence membrane charge and resistance to antimicrobial peptides (Simpson and Trent 2019). Importantly, this chemotype-level insight complements the selection analyses of HepI and OpsX by linking evolutionary constraint at the enzyme level with potential phenotypic outcomes at the membrane level. The modest relaxation of constraint observed in OpsX, particularly in regions implicated in substrate interaction, may facilitate compatibility with diverse lipid A precursors or modification states, thereby enabling flexibility in LPS assembly. When viewed alongside earlier reports of dual heptosyltransferase I systems enabling alternative inner core glycoforms, these findings collectively point toward a modular evolutionary strategy in LPS biosynthesis—where conserved catalytic cores are maintained, but peripheral components, including lipid A structure, are tuned in a lineage- and context-dependent manner.

## Conclusion

Functional studies further showed that different glycoforms can support survival, implying that duplicative retention and substrate specialization of heptosyltransferase I variants may provide ecological flexibility rather than simple redundancy. Such cases suggest that diversification of the earliest heptosylation step may be an underappreciated route to LPS heterogeneity and niche adaptation. Our analyses show that HepI and OpsX constitute functionally coupled yet evolutionarily asymmetric components of LPS inner-core biosynthesis in Gram-negative bacteria. OpsX is characterized by relaxed selective constraints and a diffuse pattern of horizontal gene transfer, whereas HepI exhibits intensified purifying selection and ecologically structured transfer. Their operon architectures further indicate pathway modularization, with HepI embedded within conserved downstream assembly modules and OpsX retaining flexibility within upstream precursor biosynthesis networks. Despite these insights, several limitations remain. First, the current study is primarily based on genomic and evolutionary inference, and therefore lacks direct biochemical validation across diverse taxa beyond previously characterized model systems. Second, functional annotation of OpsX remains incomplete in many lineages, and substrate specificity cannot be uniformly inferred from sequence or context alone. Third, ecological associations, although statistically supported, are constrained by uneven representation of environmental metadata across bacterial genomes, which may bias interpretations of lifestyle associations. Finally, the evolutionary transitions between HepI-only, OpsX-only, and dual systems remain inferential, and the temporal dynamics of these transitions cannot be directly resolved from the current dataset. Future work should focus on experimentally validating OpsX and HepI activity across representative phylogenetic groups, particularly in underexplored non-pathogenic and environmentally diverse taxa. Notably, high-resolution structural information for OpsX remains lacking, limiting detailed understanding of its catalytic architecture and substrate recognition relative to canonical WaaC-like HepI enzymes.

In addition, population-level and experimental evolution approaches could clarify whether dual heptosyltransferase I architectures provide adaptive advantages under fluctuating environmental or host-imposed pressures. Beyond this, population genomics frameworks incorporating allele frequency variation, demographic history, and population structure across ecological gradients could further test the extent of niche specialization associated with HepI and OpsX distribution patterns. Such analyses would enable a finer resolution of how selection and drift shape the maintenance, turnover, or loss of alternative LPS biosynthetic architectures in natural populations. Importantly, taxa co-encoding both HepI and OpsX represent a particularly promising system for future investigation, as they provide a natural framework to directly assess functional partitioning, regulatory dynamics, and adaptive trade-offs between paralogous pathways. Integrating transcriptomic and regulatory data may further reveal how operon modularization and gene regulation contribute to the maintenance or loss of alternative LPS biosynthetic pathways.

## Methods

### Data Retrieval and Curation from Annotree

HepI and OpsX, two variants of heptosyltransferase I involved in lipopolysaccharide (LPS) core biosynthesis, were systematically retrieved from the Annotree r214 database (Mendler et al. 2019) using their corresponding KEGG Orthology identifiers (Kanehisa et al. 2012), K02841 and K12982, respectively. The resulting datasets, comprised Gram-negative bacterial genomes in which these KO terms were annotated, along with associated metadata including genome identifiers, locus tags, genomic coordinates, and GTDB-based taxonomic classifications (Parks et al. 2022). To ensure a curated and phylogenetically representative dataset, uncultured (metagenome-assembled) genomes were excluded, type strains were removed, duplicated sequences were discarded, and only a single representative species per genus (and higher taxonomic ranks where applicable) was retained across diverse Gram-negative bacterial phyla. To construct sequence datasets suitable for downstream comparative and evolutionary analyses, coding sequences (CDS) and corresponding protein sequences were extracted by mapping Annotree-derived locus information to GTDB/RefSeq genome assemblies. Both nucleotide and amino acid datasets were retained to enable complementary analyses, including phylogenetic reconstruction and codon-aware evolutionary inference. Sequence headers were standardized to ensure traceability across datasets and integration with taxonomic metadata. Quality control steps included removal of truncated sequences (<50% of median length), dereplication of identical sequences using CD-HIT, and verification of conserved domain architecture through profile-based searches, ensuring high-confidence homolog sets.

### Selection analysis

Selection analyses were conducted to quantify differences in evolutionary constraint between HepI and OpsX across complementary phylogenetic frameworks. Codon-based multiple sequence alignments and corresponding maximum-likelihood gene trees were generated for each gene and used consistently across all downstream analyses to ensure concordance between sequence evolution and phylogenetic reconstruction. To evaluate genome-wide patterns of molecular evolution, overall dN/dS (ω) distributions were estimated for each gene using codon substitution models under the M0 (one-ratio) framework implemented in PAML codeml (Yang 2007), applied to a stratified subsampling scheme in which aligned sequences were randomly partitioned into multiple non-overlapping blocks (∼200 taxa per set) to reduce sampling bias and assess robustness of ω estimates across replicate datasets. Resulting ω distributions were summarized using parametric and non-parametric statistics, and alternative distributional fits (normal, gamma, and lognormal) were evaluated using maximum-likelihood and AIC-based model selection to characterize global selective regimes. To further resolve site-specific patterns of constraint, residue-level selection was inferred using a Bayesian framework implemented in FUBAR (Murrell et al. 2013), from which codon-wise substitution parameters (α, β) and posterior probabilities of positive selection were extracted and mapped onto amino acid positions using codon-aware alignment indexing. Residue-level ω estimates were aggregated across sites and structurally contextualized to identify conserved catalytic cores and spatial heterogeneity in selective constraint. For direct gene-to-gene comparison within identical genomic backgrounds, pairwise dN/dS values were estimated using the yn00 module (Yang 2007), restricted to taxa co-encoding both HepI and OpsX identified from AnnoTree-derived genome annotations. Average pairwise ω values were computed per species and compared between genes using matched phylogenetic ordering, and statistical significance of paired differences was assessed using non-parametric Wilcoxon signed-rank tests (Wilcoxon 1945). Across all analyses, OpsX consistently exhibited elevated ω relative to HepI, indicating comparatively relaxed purifying selection or accelerated evolutionary rates, while HepI retained stronger constraint, particularly within conserved catalytic motifs. Structural mapping of site-wise constraint was performed by projecting normalized ω values onto the three-dimensional structure of HepI in PyMOL (Schrödinger, LLC), enabling visualization of differential constraint landscapes across conserved and variable regions. Together, these complementary analyses provide concordant evidence for systematic divergence in evolutionary constraint between HepI and OpsX across global, site-specific, and paired genomic contexts.

### RELAX analysis of selection intensity

Selection intensity differences between host-associated and non-host-associated lineages were quantified using the RELAX framework implemented in HyPhy (Wertheim et al. 2015), which estimates shifts in the distribution of dN/dS (ω) through the selection intensity parameter (K). For both HepI and OpsX, codon-based multiple sequence alignments and corresponding codon-informed maximum-likelihood gene trees were used to ensure consistency between sequence evolution and phylogenetic inference. Lifestyle metadata were systematically derived from genome-associated records obtained from the BV-BRC (formerly PATRIC) (Davis et al. 2020), where species were classified into ecological categories (Pathogen, Opportunist, Non-pathogen) based on aggregated host, isolation, and clinical annotations. In the global dataset, host-associated lineages were defined stringently by designating only Pathogen taxa as the foreground and Non-pathogen taxa as the background to maximize ecological contrast and reduce ambiguity arising from intermediate lifestyles; alignments were filtered accordingly and gene trees were pruned to match the retained taxa. Terminal branches were labeled according to lifestyle assignments, while internal branches were conservatively assigned to the background class. RELAX was executed under a discrete three-rate ω distribution using the “Minimal” model, with a minimum of 30 taxa per category enforced to ensure statistical robustness, and likelihood ratio tests (LRTs) were used to evaluate deviations from the null hypothesis (K = 1). In the coexisting dataset, restricted to taxa encoding both HepI and OpsX, the same pipeline was applied; however, Pathogen and Opportunist categories were combined to define a broader host-associated foreground, while Non-pathogen taxa served as the background, reflecting a continuous spectrum of host association and ensuring adequate foreground representation under reduced sample size, without imposing strict minimum thresholds. Across both datasets, selection intensity (K) and associated p-values were extracted and visualized, with K > 1 indicating intensified selection and K < 1 indicating relaxation, thereby enabling direct comparison of evolutionary constraints across genes and ecological contexts.

#### Comparative Phylogenetic Structure

Phylogenetic structure across genomes co-encoding HepI and OpsX was examined using a controlled subset of taxa in which both genes are present, enabling direct comparison under a shared genomic background. Protein sequences for each gene were aligned independently using MAFFT (Katoh & Standley 2013) with automatic strategy selection to ensure alignment accuracy across diverse bacterial lineages. Maximum likelihood phylogenies were then reconstructed using IQ-TREE (v2) (Minh et al. 2020) with ModelFinder-based substitution model selection (MFP) (Kalyaanamoorthy et al. 2017), and branch support was assessed using 1000 ultrafast bootstrap and 1000 SH-aLRT replicates (Hoang et al. 2018; Guindon et al. 2010). To maintain consistency in evolutionary orientation, *HepIII* homologs from *Escherichia coli* were used as a fixed outgroup for both gene trees. Resulting trees were processed and visualized using ETE3 (Huerta-Cepas et al. 2016), where trees were rooted using the specified outgroup, ladderized, and branch lengths optionally normalized by total tree depth to facilitate cross-gene comparison. Taxonomic metadata were incorporated to annotate leaves based on ecological categories (pathogen, opportunist, non-pathogen), with labels standardized to genus–species format and rendered using color-coded node styles; individual trees were exported at high resolution and combined into a single comparative figure using the Python Imaging Library. To quantitatively assess evolutionary divergence patterns, branch lengths were extracted from the rooted trees and classified as either tip or internal branches. Statistical comparisons of branch length distributions were performed both between genes (HepI vs OpsX for each branch type) and within genes (internal vs tip) using the non-parametric Mann–Whitney U test implemented in SciPy (Virtanen et al. 2020), with summary statistics and p-values recorded for reproducibility. Branch length distributions were visualized using boxplots and swarm plots generated with Seaborn (Waskom 2021), alongside heatmaps summarizing mean branch lengths across categories. All analyses were conducted using reproducible Python scripts with standardized input formats, ensuring methodological consistency and minimizing biases associated with phylogenetic reconstruction and comparative inference.

### Gene–Species Tree Reconciliation and Horizontal Gene Transfer Inference

Gene–species tree reconciliation was performed to infer duplication, transfer, and loss (DTL) events for *hepI* and *opsX* across global and coexisting datasets. For the global dataset, protein sequences were clustered at 95% identity using CD-HIT (Li & Godzik 2006) to reduce redundancy, followed by multiple sequence alignment with MAFFT (Katoh & Standley 2013). Alignments were subsequently filtered to remove gap-rich sequences and trimmed based on column occupancy to retain phylogenetically informative regions. Maximum likelihood gene trees were inferred using IQ-TREE (Minh et al. 2020) with bootstrap support, and poorly supported nodes (<70%) were collapsed into polytomies using the R package ape. Trees were midpoint-rooted and reconciled against a curated bacterial reference species tree (bac120) after taxon mapping (Parks et al. 2018; Parks et al. 2022). Reconciliation analysis was conducted using RANGER-DTL (Bansal et al. 2012) under a duplication–transfer–loss framework, with 50 independent runs performed to account for stochastic variation. Event outputs were aggregated to estimate frequencies and support values for inferred events, and high-confidence transfers were identified based on recurrence across runs. For the coexisting dataset, gene trees for *hepI* and *opsX* were directly reconciled against the same species tree without clustering or extensive preprocessing. Post-analysis included aggregation of event frequencies and construction of transfer networks, with additional integration of ecological metadata (pathogen, non-pathogen, opportunist) from BV-BRC (Davis et al. 2020) to assess transfer patterns across lifestyle categories.

### Codon Usage Bias Analysis

Codon usage bias was assessed for genes of interest using the subset of bacterial genomes co-encoding both *hepI* and *opsX*, enabling direct comparison within shared genomic backgrounds. Coding sequences (CDS) for each gene were extracted and analyzed using a customized Python-based workflow implemented with Biopython, NumPy, pandas, and SciPy. Multiple complementary metrics were employed to capture distinct aspects of codon usage bias. GC content at third codon positions (GC3) was calculated for each coding sequence as a proxy for nucleotide compositional bias (Sharp & Li 1986). The effective number of codons (ENC) was estimated using the Wright approximation to quantify overall codon usage uniformity independent of amino acid composition (Wright 1990). Relative synonymous codon usage (RSCU) values were computed for each codon across sequences (Sharp & Li 1987), and mean codon usage profiles were compared between gene sets, with per-codon statistical testing performed using paired comparisons. In addition, codon adaptation index (CAI) values were calculated to assess translational optimization (Sharp & Li 1987), using organism-specific reference sets derived from highly expressed housekeeping genes (including ribosomal proteins, elongation factors, and RNA polymerase subunits) retrieved from public databases. Codon weights for CAI were computed from aggregated reference sequences and applied to individual CDS.

For all metrics, distributions were first evaluated for normality using the Shapiro–Wilk test, and appropriate statistical comparisons were performed using either parametric (Student’s or Welch’s t-test) or non-parametric (Mann–Whitney U or Wilcoxon signed-rank test) approaches as dictated by data properties (Shapiro & Wilk 1965; Student 1908; Welch 1947; Mann & Whitney 1947; Wilcoxon 1945). Summary statistics, including means, variances, and standard deviations, were computed for each gene set, and effect sizes were estimated where appropriate. Codon usage distributions and comparative patterns were visualized using publication-quality plots, including density distributions, boxplots, violin plots, and codon-level bar representations. This multi-metric framework follows established practices in codon usage analysis and enables robust characterization of compositional bias, codon preference, and translational adaptation across gene sets within comparable genomic contexts.

### Synteny analysis

Synteny analysis was performed to investigate the genomic neighborhood organization of *hepI* and *opsX* across bacterial genomes using a customized Python-based pipeline. Coding sequences were screened for target genes based on curated annotations, and local genomic neighborhoods were extracted within a ±5 kb window to capture operon-scale gene context. Gene annotations were standardized using a rule-based framework to harmonize synonymous functional labels, enabling consistent cross-genome comparison. A genome-by-gene presence–absence matrix was constructed to quantify conservation of neighborhood components, and hierarchical clustering based on compositional similarity was used to compare syntenic architectures across genomes. Putative operon structures were inferred from conserved gene order within each neighborhood, and recurrent architectures were identified based on frequency across genomes. Gene neighborhood and operon structures were visualized using established bacterial synteny frameworks in a clinker/EasyFig-like style (Gilchrist & Chooi 2021; Sullivan et al. 2011), with all analyses implemented in Python. The workflow is conceptually consistent with gene neighborhood–based synteny and pan-genome approaches commonly used in microbial comparative genomics. The same pipeline was applied to the subset of taxa co-encoding *hepI* and *opsX*, which were analyzed separately to resolve syntenic organization under shared genomic contexts.

### Mobile Genetic Element Landscape Profiling Across Genomes

Genomic context analysis was performed in conjunction with synteny reconstruction to characterize the local genomic environments of *hepI* and *opsX* associated loci across two defined datasets: a global dataset comprising all bacterial genomes encoding either gene system to capture broad phylogenetic diversity, and a coexisting dataset restricted to genomes harboring both systems to enable context comparison within shared genomic backgrounds. Gene neighborhoods were extracted using a customized python-based pipeline and analyzed at two spatial resolutions, containing a ±5 kb window to resolve operon-scale syntenic organization and a ±10 kb window to evaluate broader genomic island–scale contextual signals. Coding sequences were identified based on curated genome annotations, and functional descriptors were standardized using a rule-based harmonization framework to ensure consistent cross-genome comparison of gene content and neighborhood architecture. Putative mobile genetic elements (MGEs) were identified using a functional signature–based heuristic screening strategy, whereby genes were classified based on curated mobility-associated annotations, including insertion sequences and transposases (IS families), integrases, recombinases, prophage-associated structural genes, conjugation machinery (type IV secretion system components and VirB proteins), relaxases, and plasmid-associated replication and transfer genes. This gene-content–based screening was applied as a rapid mobilome proxy for large-scale comparative genomic analysis. Genomic proximity between loci of interest and MGE-associated features was computed using coordinate-based distance estimation, with overlapping features assigned a distance of zero and loci within defined thresholds considered part of extended genomic neighborhoods. Species-level presence–absence matrices of MGE signatures were constructed and subjected to hierarchical clustering to compare genomic neighborhood composition across taxa. The overall analytical framework follows established practices in microbial comparative genomics for gene neighborhood and mobilome assessment and is broadly consistent with gene neighborhood–based synteny representation strategies and genome-scale mobilome inference frameworks commonly used for contextual interpretation of horizontally acquired genomic regions.

## Author contributions

A.M.G, P.A. and E.A.T were all involved in the Conceptualization, Data curation, Formal analysis, Methodology, Validation, Visualization and Writing – review and editing; A.M.G. was responsible for the Investigation and Writing – original draft; P.A. and E.A.T were involved in the Project administration and Supervision

