## Supplemental for "HepI and OpsX are functionally coupled but evolutionarily asymmetric heptosyltransferase variants: ecological transitions and operon modularization drive divergent constraints and flexibility"

Figure legends

**SI Fig. S1.** (a) Heatmap of mean branch lengths showing higher internal and tip divergence in HepI compared to OpsX, with greater separation between branch types in HepI. (b) Distribution of predicted lipid A chemotypes across coexisting taxa, highlighting a predominance of the tetra-acyl | Kdo₂ | PEtN form over hexa-acylated variants.

**SI Fig. S2.** (a) HepI shows a strong lifestyle-associated HGT signal (β = 3.88, p < 10⁻¹⁰; permutation p = 0.014) with enrichment on internal branches (~55%). (b) OpsX shows no significant association (β = 0.57, p = 0.51) and is dominated by terminal transfers (~52%). (c) Despite comparable HGT counts (HepI n = 20; OpsX n = 24), transfer dynamics differ markedly, with HepI reflecting deeper, ecologically structured integration and OpsX showing predominantly recent, transient acquisitions.

**SI Fig. S3.** Phylogenetic reconciliation shows HepI transfers (orange; branch width indicates confidence) are clustered within ecologically coherent clades, whereas OpsX transfers are scattered and diffuse.

**SI Fig. S4.** Codon usage bias analysis of co-encoded genes shows no significant differences between HepI (steel blue) and OpsX (tomato red) across (a-d) GC3, ENC, RSCU, and CAI metrics, indicating comparable mutational and translational constraints.

**SI Fig. S5.** Genomic context analysis shows no consistent enrichment of mobile genetic element signatures around HepI (a-global; c-coexisting) or OpsX loci (b-global; d-coexisting). Occasional MGE-associated genes are rare, sporadic, and non-recurrent, with no locus- or lineage-specific pattern, indicating that both genes are shaped by long-term genomic remodeling rather than recent MGE-driven acquisition.

Fig S1

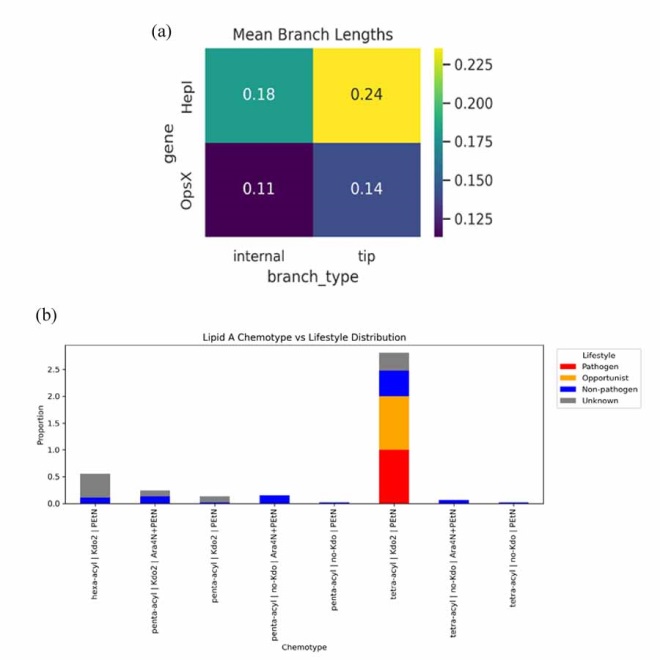

Fig S2

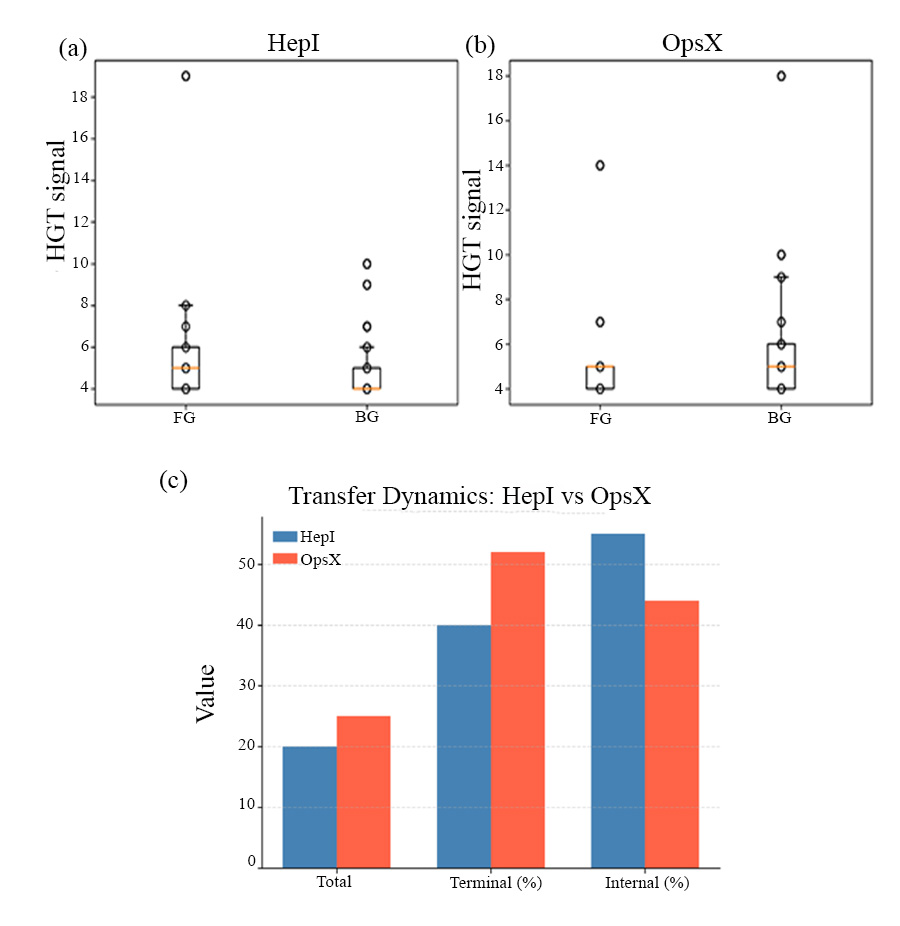

Fig S3

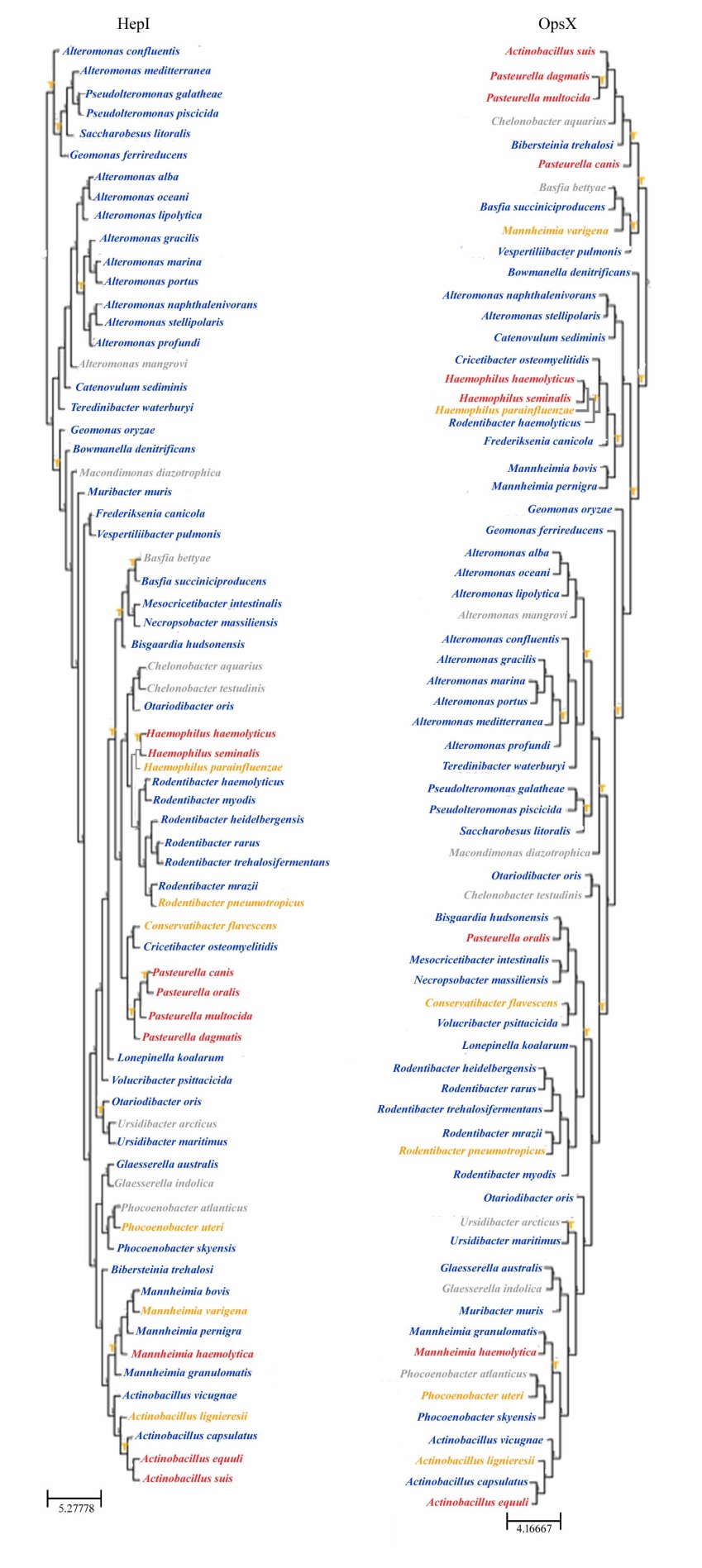

Fig S4

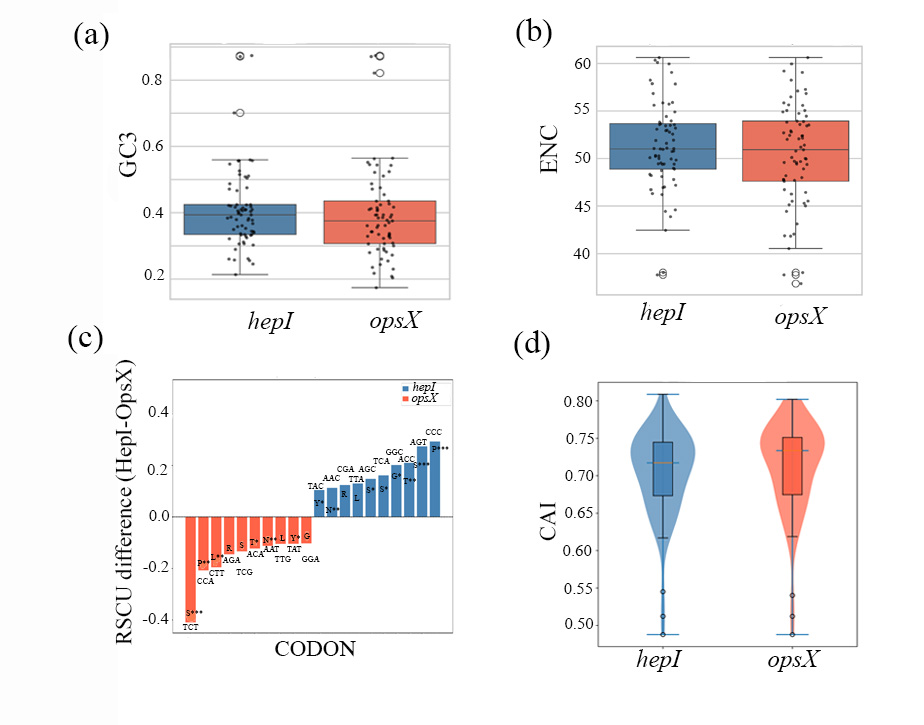

Fig S5

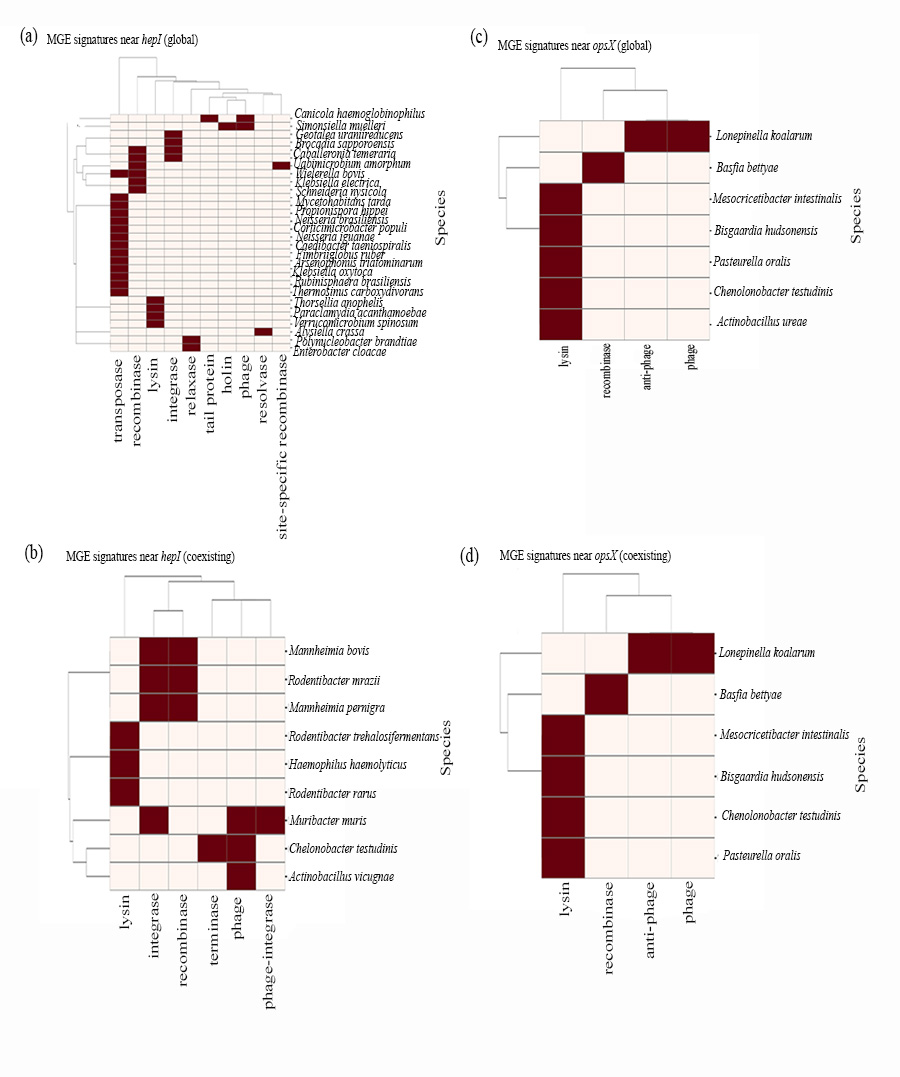

Table S4

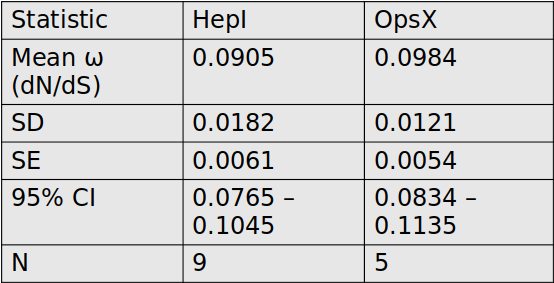
